# Graph theoretic analysis reveals intranasal oxytocin induced network changes over frontal regions

**DOI:** 10.1101/2020.11.22.393561

**Authors:** Shuhan Zheng, Diksha Punia, Haiyan Wu, Quanying Liu

**Affiliations:** Southern University of Science and Technology, Shenzhen 518005, China; Department of Psychology, University of Macau, Macau, China; University of California, Irvine, USA

**Keywords:** graph theory, fMRI, oxytocin, frontal cortex, Interpersonal Reactivity Index

## Abstract

In this study, we aim to elucidate how intranasal oxytocin modulates brain network characteristics, especially over the frontal network. As an essential brain hub of social cognition and emotion regulation, we will also explore the association between graphic properties of the frontal network and individual personality traits under oxytocin (OT) administration. 59 male participants administered intranasal OT or placebo were followed by restingstate fMRI scanning. The Correlation-based network model was applied to study OT modulation effects. We performed community detection algorithms and conducted further network analyses, including clustering coefficient, average shortest path and eigenvector centrality. In addition, we conducted a correlation analysis between clustering coefficients and the self-assessed psychological scales. Modular organizations in the OT group reveal integrations of the frontoparietal network (FPN) and the default mode network (DMN) over frontal regions. Results show that frontal nodes within the FPN are characterized by lower clustering coefficients and higher average shortest path values compared to the placebo group. Notably, these modulation effects on frontal network property are associated with Interpersonal Reactivity Index (IRI) fantasy value. Our results suggest that OT elevates integrations between FPN, DMN and limbic system as well as reduces small-worldness within the FPN. Our results support graph theoretic analysis as a potential tool to assess OT induced effects on the information integration in the frontal network.

## Introduction

Oxytocin (OT) is a neuropeptide closely linked to social adaptation and prosocial behaviors (Churchland and Winkielman 2012; Jones et al. 2017; Y. Ma et al. 2016; Ross et al. 2009). Previous work has suggested that OT promotes social cooperation (Zak et al. 2007) and trust (Kosfeld et al. 2005). Emerging evidence has shown that OT can enhance motivation or sensitivity to social cues (Bartz et al. 2010; Hurlemann et al. 2010; Neumann 2008), e.g., emotion recognition (Domes et al. 2007), and in-group and out-group processes (De Dreu et al. 2011). However, a recent replication study reported no effect of OT administration on trust behaviors (Declerck et al. 2020). The effects of OT on social behaviors are still debated. A growing need for accurate applications of OT in multiple fields, including treatment for social deficits in neurodevelopmental disorders and explanatory factors for individual social difference (Baribeau and Anagnostou 2015; Parker et al. 2017), calls for a comprehensive understanding of both the behavioral effect of OT and its underlying neural basis.

Following the line of behavioral research, the mechanism of the effects of OT have been studied at different levels, from microscopic molecules to macroscopic brain networks. At the molecular level, OT receptors have been found in the hippocampus, cingulate cortex, amygdala and particularly in the medial prefrontal cortex (Boccia et al. 2013; Raam 2020).

At the macroscopic level, many previous studies have reported that OT alters functional connectivity and neural activation across multiple brain areas (Bethlehem et al. 2017; Riem, Bakermans-Kranenburg, Pieper, et al. 2011; Z. Zhao et al. 2019), including the amygdala, hippocampus, medial frontal cortex, orbitofrontal cortex, and anterior cingulate cortex (Eckstein et al. 2017; Riem, Van Ijzendoorn, et al. 2012; Sripada et al. 2013).

Resting-state fMRI, as a non-invasive way to measure the spontaneous neural activity in the human brain, has been applied to study differential pharmacological effects on brain functional interaction in individuals (see a recent review (Seeley et al. 2018)). Researchers have gained insight into the effects of OT administration on human brain function, especially in the social brain network, from both healthy and clinical samples (Frijling et al. 2016; Watanabe et al. 2015). However, rather than a widely distributed impact on various brain regions, the modulation effects of OT are considered mainly focused on the amygdala-related circuitry, the functional connectivity involved in the amygdala region (Dodhia et al. 2014; Eckstein et al. 2017; Koch et al. 2016), and the mentalizing related functional connectivity of the temporo-parietal junction (TPJ) (Wu et al. 2020). Additionally, within its role in the default mode network (DMN) (Mars et al. 2012; Xie et al. 2016), one study found OT effectively altered connectivity within the DMN (Kumar et al. 2019).

One possible mechanism underlying the effects of OT on affiliative and prosocial behaviors is that OT selectively modulates brain circuits involved in social processing. Although the amygdala is the most used seed in seed-based connectivity analysis, it is important to note that OT mediates amygdala connectivity with other brain regions through the frontal network (Sripada et al. 2013) (e.g., the pregenual anterior cingulate cortex (pgACC)) (Y. Fan, Herrera-Melendez, et al. 2014). As proposed by Quintana and colleagues, one central route to the brainstem by intranasal OT administration is via the trigeminal nerve, to the amygdala and the prefrontal cortex (PFC) via the olfactory bulb (Quintana et al. 2015). This leads to our hypothesis that intranasal OT changes the network characteristics over the frontal brain network.

Previous research has reported the ventromedial prefrontal cortex (vmPFC) plays as a hub in the generation of affective meaning, bridging conceptual and affective processes (Roy et al. 2012). Evidence has also shown that OT alters the response of the inferior frontal gyrus when exposed to infant crying (Riem, Bakermans-Kranenburg, Pieper, et al. 2011; Riem, Bakermans-Kranenburg, and van IJzendoorn 2016). The dorsomedial prefrontal cortex (DMPFC) and dorsal anterior cingulate cortex (dACC) form emotional responses to other people’s mental states (Y. Fan, Niall W Duncan, et al. 2011a). The medial prefrontal cortex is critical in emotion regulation, and has been reported to modulate amygdala output to assign awareness of emotional visual stimuli (Banks et al. 2007; Mitchell and Greening 2012; Morawetz et al. 2017). Since the frontal network overlaps with the social brain network and plays a critical role among different subnetworks (Cole et al. 2014; Li et al. 2014; Mars et al. 2012), it is important to understand how OT modulates different subnetworks through frontal regions.

As a novel brain network model, graph-based network analyses are capable of uncovering system-level alterations associated with different processes in the resting brain without losing too many biological details (Jinhui Wang et al. 2010). In this work, we applied the network model to study the modulation effects of OT over the frontal network by means of a multi-method brain network approach combining placebo-controlled intranasal OT administration.

We investigated the modular structure using community detection algorithms, which revealed the modular structure through maximized network modularity. However, community detection is not capable of uncovering microscopic details, which strongly limit its efficacy in studying the property of a single node. Therefore, we applied other network algorithms, including a clustering coefficient (CC), average shortest path (ASP) and eigenvector centrality (EC), all allowing us to reveal the property of a node locally.

The clustering coefficient is a critical measurement of the local connectivity in the brain network model (Stam 2014). Many studies use the clustering coefficient as a measure to reflect brain topology. Moreover, it has been widely shown that the clustering coefficient is related to cognitive performance and mental disease (Langer et al. 2012; X. Zhao et al. 2012). The average shortest path of the brain network represents the capability of the brain to integrate information and it is a commonly used indicator for the small-worldness (Karwowski et al. 2019; Stam 2014). The eigenvector centrality describes a node’s influence on the brain network. As a node has higher EC value, the node has more important neighbour nodes, but does not necessarily have more edges connecting other nodes. This property permits EC to be an important measure in the brain network framework (Lohmann et al. 2010). Recent works have shown the eigenvector centrality is a potential biomarker for Alzheimer’s disease as well as diabetes, and can be used to highlight applicable targets for brain stimulation (Antonenko et al. 2018; Binnewijzend et al. 2014; van Duinkerken et al. 2017). The graphs of theoretical measures, together, aim to comprehensively examine the characteristics of brain network property under OT administration.

Neuroscience has built an account of the effects of OT on social behaviors and causes of change at the neuronal level. We aim to further focus on how OT effects the frontal network, and how those effects are linked to social personality traits.

The Interpersonal Reactivity Index (IRI) quantifies individual cognitive and emotional reactions to a wide range of interpersonal events and stimuli (Davis et al. 1980; Keaton 2017). The IRI fantasy assesses the extent to which individuals imaginatively transpose themselves into fictional situations. In this aspect, to transpose requires an ability to break through the self-other boundary. Not only is this related to immersing oneself into anothers mental state, but it is also related to vicarious emotions/experience by way of putting oneself into a movie or in a novel characters shoes and so on. Task-based fMRI studies have demonstrated prefrontal engagement during action or intention imagination tasks (Macuga and Frey 2012)and the ventromedial prefrontal cortex (vmPFC) showed overlapping responses to personal and vicarious rewards (Mobbs et al. 2009). Furthermore, meta-analyses of the empathy neural network have identified a high probability of activation over the frontal area, including dACC, aMCC-SMA (Y. Fan, Niall W. Duncan, et al. 2011b; Jauniaux et al. 2019). There is also a “gate theory” hypothesis of the frontal area, with evidence showing that oxytocin-sensitive neurons in the prefrontal cortex act as a “gate” in social memory (Raam 2020), as it represents an instance of when an individual imagines a stimulus, rather than sees it. Recent studies have shown the association between spontaneous mind-wandering and creativity in resting states with subnetworks (FPN and DMN) property (Golchert et al. 2017; Shi et al. 2018). These findings together lead to potential theories related to the frontal network with the degree of spontaneous mind-wandering, social imagination and creativity during a resting state, and how OT alters the related states. Previous research has found individuals with low score in IRI empathy show greater improvement in the Reading the Mind in the Eyes Test (RMET) performance after OT administration compared to placebo (Radke and de Bruijn 2015) however, the study lacks further investigation of underlying network alterations associated with RMET performance. It does show that IRI subscales can be a window to investigate OT modulation effects. Based on previous literature about relations between individual fantasy level and the frontal network, together with relations between the frontal network and OT, we hypothesized the following potential brain network effects about enhanced social function after OT administration: 1) OT can increase the integration between different subnetworks throughout frontal regions; 2) locally, OT can reduce the small-worldness property within the FPN; 3) from the individual perspective, OT has more substantial modulation effects on the FPN for individuals with lower fantasy trait scores.

## Methods

To test these hypotheses, we performed analysis as shown in the flow diagram **Figure 1**. We recruited 59 healthy male participants, and randomly assigned them to two groups (an oxytocin group and placebo group). After the standard fMRI preprocessing, we applied connectivity analysis on the BOLD signals averaged across voxels in 90 AAL regions. Then, we performed community detection to characterize network structural differences between the two groups. Next, we applied local network measures, such as CC and ASP, to identify node-wise variability. Finally, we conducted correlation analyses to associate the graph properties with personal traits. All connectivity analyses for network constructions and graph theoretical analyses were done in Matlab 2019b (The Mathworks Inc., Natick, MA, USA). Correlation analyses were performed in R.

**Figure 1:**
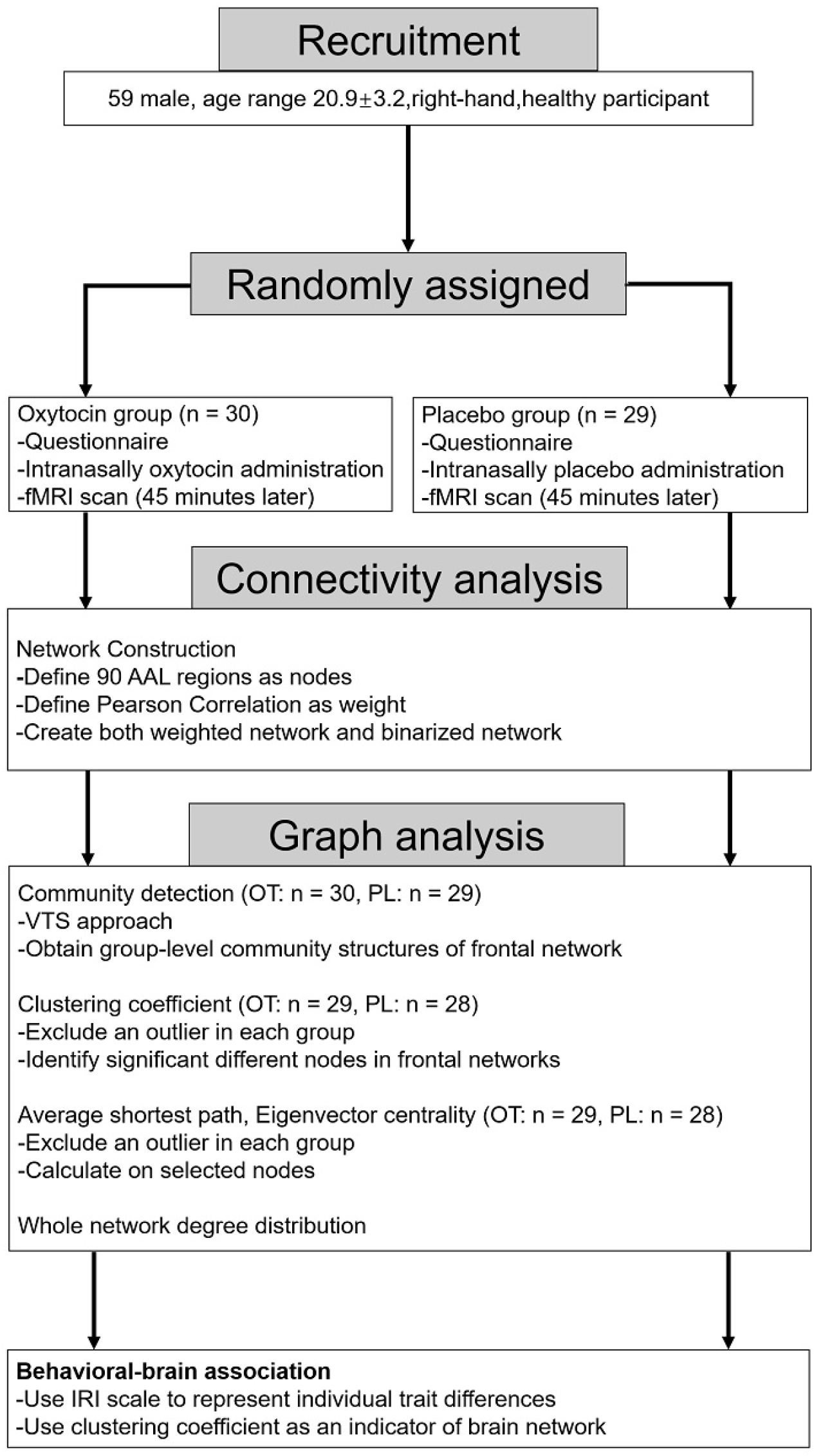
Analysis flow diagram. A brief description is provided in each box, summarizing key points in each analysis step.

### Participants

As many reports have pointed out, OT’s effects on brain connectivity could be affected by age, sex, and level of proficiency (Bartz et al. 2010; Campbell et al. 2014; Ebner et al. 2016; Luo et al. 2017). To avoid the effects of these confounders, we recruited 59 healthy male participants who were 20.9 ±2.32 years of age, were righthanded, and had 13~18 years of education. These individuals participated in this study via an online recruiting system. All participants filled out a screening form, and participants were included in the study only if they confirmed they were not suffering from any significant medical or psychiatric illness, not using medication, and not drinking and/or smoking on a daily basis. Participants were instructed to refrain from smoking or drinking (except water) for 2 hours before the experiment. Participants received full debriefing upon completion of the experiment. Informed written consent was obtained from each participant before each experiment. The study was approved by the local ethics committee.

### Study Design and Drug Administration

Below, we describe the design and drug administration procedures. Here, we used a double-blind placebo-controlled group design where participants were randomly assigned to the oxytocin (OT) group or the placebo (PL) group (**Table 1**). As we reported in the previous study (Wu et al. 2020), each participant visited the lab twice. The first visit was for filling out questionnaires (e.g., the Interpersonal Reactivity Index (IRI)), (Radke and de Bruijn 2015) and the second visit was for MRI scanning.

**Table 1:** Table representation of community detection result.

| PL-C1 | OT-C1 | PL-C2 | OT-C2 | OT-C3 |
| --- | --- | --- | --- | --- |
| MFG. L&R * |  | ORBsupmed.L&R |  | IFGoperc.L&R* |
| IPL L&R |  | ORBsup.L&R |  | MTG.L&R |
| PCUN.L&R |  | OLF L&R |  | IFGtriang.R* |
| ANG.R |  | REC.L&R |  | IFGtriang.L |
| ITG.R |  | TPOmid.L&R |  | ORBinf.L&R |
| ORBmid.R* |  | ANG.L | AMYG L&R | SFGmed |
| IFGoperc.L&R* | SFG.L&R | ACG.L&R | HIP.L&R | TPOsup.R |
| IFGtriang.R* | PCG.L&R | PCG.L&R | PHG L&R | ITG.L |
| IFGtriang.L | ANG.L | ORBinf.L&R |  |  |
| SMG.L&R |  | SFGmed.L&R |  |  |
| MCG.R |  | SFG.L&R |  |  |
| INS.R |  | MTG.R |  |  |
| ITG.L |  | TPOsup.R |  |  |
| PreCG.L |  |  |  |  |
| ORBmid.L |  |  |  |  |

Each participant arrived for the second visit 1 hour before MRI scanning. At 45 minutes before the MRI scanning, a single dose of *24IU* oxytocin or placebo was administered intranasally by three puffs of 8IU per nostril. The puffs were delivered to each nostril in an alternating order and with a 45-second break between each puff to each nostril. The resting MRI scan lasted around 5 minutes and the subjects were instructed to keep their eyes closed, but to not fall sleep.

### MRI data acquisition and analysis

All images were acquired on a 3T Siemens Tim Trio scanner with a 12-channel head coil. Functional images employed a gradient-echo echo-planar imaging (EPI) sequence with the following MRI scanner parameters:(40ms TE, 2s TR, 90°flip, 210mm FOV, 128 by 128 matrix, 25 contiguous 5mm slices parallel to the hippocampus and interleaved). We also acquired whole-brain T1-weighed anatomical reference images from all participants (2.15ms TE, 1.9s TR, 9°flip, 256mm FOV, 176 sagittal slices, 1 mm slice thickness, perpendicular to the anterior-posterior commissure line).

fMRI data preprocessing was performed using Statistical Parametric Mapping software (SPM12: Wellcome Trust Centre for Neuroimaging, London, UK). The functional image time series were preprocessed to compensate for slice-dependent time shifts, motion corrected, and linearly detrended. Next the images were coregistered to the anatomical image, spatial normalized to Montreal Neurological Institute (MNI) space, (http://www.bic.mni.mcgill.ca/ServicesAtlases/HomePage) and spatially smoothed by convolution with an isotropic Gaussian kernel (full width at half maximum, FWHM=6 mm). The fMRI data were high-pass filtered with a cutoff of 0.01 Hz. The white matter (WM) signal, cerebrospinal fluid (CSF) signal and global signal, as well as the 6-dimensional head motion realignment parameters, the realignment parameters squared, their derivatives, and the squares of the derivatives were regressed. The resulting residuals were then low-pass filtered with a cutoff of 0.1 Hz.

### Network construction

After the fMRI data preprocessing, we extracted the brain signals from 90 brain regions according to the Automatic Anatomical Labeling (AAL) brain atlas. The signals from the same brain regions were averaged, and we calculated the Pearson correlation across the signals of 90 brain regions pairwise and obtained the weighted adjacency matrix.

Previous research has shown that gross inferences of graph topology, like whether the brain network was small-world or scale-free, are robust across different network densities. But both absolute values of specific parameters such as path length, clustering coefficient, and degree distribution descriptors vary considerably across network density (Fornito et al. 2010; Yu et al. 2018). Thus, in order to obtain robust results, we constructed the binary adjacency matrix by setting different edge density levels. For example, a 10% edge density level means the top 10% weighted edges were set as one while other edges were ignored as zero. Binary and weighted adjacency matrices were both used in calculations. At present, the meaning of a negative correlation is not clear in fMRI-based network construction. We set negative matrix elements as zero.

### Graph theoretical analysis

To test our hypotheses, we used multiple graph theoretical measures including community detection (CD), clustering coefficient (CC), average path length (ASP), eigenvector centrality (EC) and the degree of distribution over the whole brain network model. We applied the brain connectivity toolbox (BCT) in Matlab for the graph theoretical analysis (Rubinov and Sporns 2010).

First, we studied the network modularity through the lens of community detection. The group-level network for CD is constructed using the virtual-typical-subject (VTS) approach (Taya et al. 2016), which used group averaged data to extract patterns in the network. Then we applied the Louvain heuristics algorithm, with the structural resolution parameter *γ*= 1.2. Here, *γ* controls the modular size in the community detection result.

In the undirected network framework, clustering coefficient is defined as following.

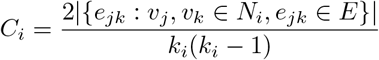

where *e_jk_* is the edge between nodes *v_j_* and *v_k_*. *N_i_* is the set of nodes directly connected to node i, and *E* is the set of edges.

We calculated ASP over the weighted undirected adjacency matrix. Typically, ASP is defined by the following formula.

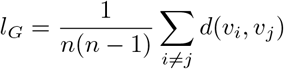

In this definition, *l_G_* is a global network property, which is not applicable in our study for we focus on the frontal area in our hypotheses. Adopting the ASP definition directly in our analysis will take other brain areas and their spurious correlations into account. Spurious correlations in ASP lead to longer path values. This entails unwanted impacts to ASP as a measure reflecting information integration in the frontal regions. Therefore, we adjusted the traditional ASP with the following procedures in this study:

1. The length between any node is 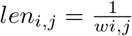
2. We calculate shortest path between any node *i* and node *j*, denoted as *d_ij_*. Note that the shortest path has a minimal weighted distance, not a minimal number of edges.
3. For node *i* ∈ frontal areas, we select node *k* satisfying *w_ik_* ≠ 0. We define *l_i_* as the mean of *d_jk_*.

OT could selectively activate or deactivate brain areas, resulting from changes in node centrality in the PL and OT networks. We applied the eigenvector centrality to assess the between-group centrality differences in the frontal area. To uncover topological property in the brain network of two groups, we examined the degree of distribution on the whole brain network level.

## Results

### Community detection

The maximum Q value of detected modules, which quantified the modularity of corresponding networks,is 0.259 for OT and 0.262 for the PL group. The maximum Q value is obtained from applying the community detection method in binary networks with an edge density level of 4%. As shown in **Figure 2**, the frontal network of the OT group exhibited more modules than the PL group. Detailed community detection result are listed in **Table 1**. The correspondence between full names and abbreviations are listed in **Table 2**. In **Table 1**, nodes indicated with a star are ROIs where clustering coefficients are significantly different between OT and PL groups. Shared nodes were listed in merged cells. Amygdala (AMYG), hippocampus (HIP) and parahippocampal gyrus (PHG) were not incorporated in modules of the PL group. While in the OT group, AMYG, HIP and PHG were all components of the OT-C2 module, together with nodes in the orbitofrontal cortex.

**Figure 2:**
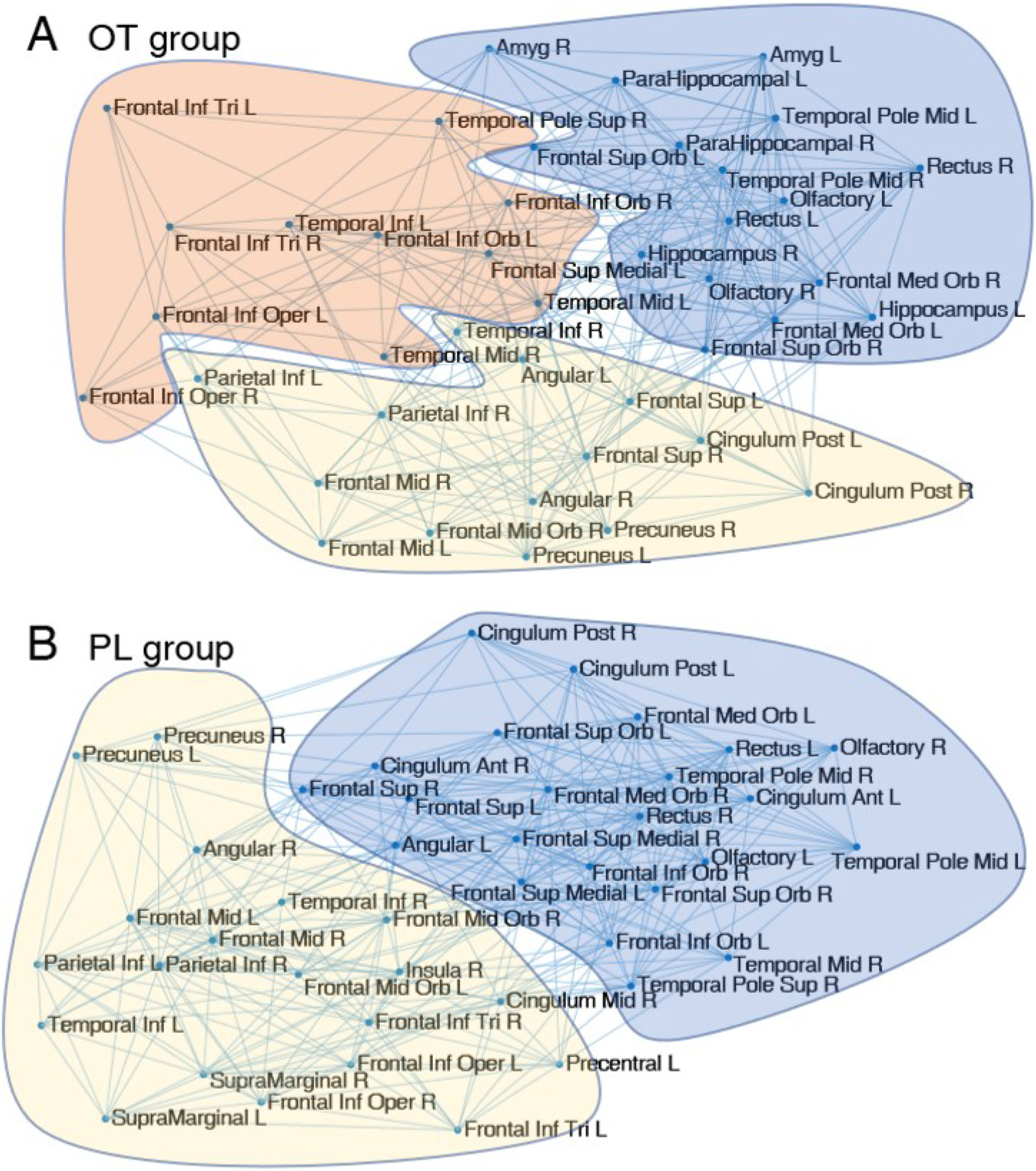
Community detection result. (A) In the OT group, nodes in the frontal area were found and classified into 2 modules, including other highly connected nodes. (B) PL group reflected a more segregated pattern in the frontal network and highly connected nodes. The same colour suggests the corresponding 2 modules have the maximum number of the same nodes.

**Table 2:** Brain regions with abbreviations and MNI coordinates

| Region name | Abbreviation | MNI Coord. | Region name | Abbreviation | MNI Coord. |
| --- | --- | --- | --- | --- | --- |
| Frontal Mid L | MFG.L | -33, 33, 35 | Amygdala R | AMYG.R | 27, 2, -18 |
| Frontal Mid R | MFG.R | 38, 33, 34 | Hippocampus L | HIP.L | -25, -21, -10 |
| Parietal Inf L | IPL.L | -43, -46, 47 | Hippocampus R | HIP.R | 29, -20, -10 |
| Parietal Inf R | IPL.R | 46, -46, 50 | ParaHippocampal L | PHG.L | -21, -16, -21 |
| Precuneus L | PCUN.L | -7, -56, 48 | ParaHippocampal R | PHG.R | 25, -15, -20 |
| Precuneus R | PCUN.R | 10, -56, 44 | Temporal Inf L | ITG.L | -50, -28, -23 |
| Angular R | ANG.R | 46, -60, 39 | Temporal Inf R | ITG.R | 54, -31, -22 |
| Angular L | ANG.L | -44, -61, 36 | Temporal Mid L | MTG.L | -56, -34, -2 |
| Frontal Mid Orb L | ORBmid.L | -31, 50, -10 | Temporal Mid R | MTG.R | 57, -37, -1 |
| Frontal Mid Orb R | ORBmid.R | 33, 53, -11 | Temporal Pole Mid L | TPOmid.L | -36, 15, -34 |
| Frontal Inf Oper L | IFGoperc.L | -48, 13, 19 | Temporal Pole Mid R | TPOmid.R | 44, 15, -32 |
| Frontal Inf Oper R | IFGoperc.R | 50, 15, 21 | SupraMarginal L | SMG.L | -56, -34, 30 |
| Frontal Sup Orb L | ORBsup.L | -17, 47, -13 | SupraMarginal R | SMG.R | 58, -32, 34 |
| Frontal Sup Orb R | ORBsup.R | 18, 48, -14 | Cingulum Post L | PCG.L | -5, -43, 25 |
| Frontal Med Orb L | ORBsupmed.L | -5, 54, -7 | Cingulum Post R | PCG.R | 7, -42, 22 |
| Frontal Med Orb R | ORBsupmed.R | 9, 51, 30 | Cingulum Ant R | ACG.R | 8, 37, 16 |
| Frontal Inf Tri L | IFGtriang.L | -46, 30, 14 | Cingulum Ant L | ACG.L | -4, 35, 14 |
| Frontal Inf Tri R | IFGtriang.R | 50, 30, 14 | Cingulum Mid R | MCG.R | 8, -9, 40] |
| Frontal Sup L | SFG.L | -18, 35, 42 | Olfactory L | OLF.L | -8, 15, -11 |
| Frontal Sup R | SFG.R | 22, 31, 44 | Olfactory R | OLF.R | 10, 16, -11 |
| Frontal Inf Orb L | ORBinf.L | -36, 31, -12 | Insula R | INS.R | 43, 19, -5 |
| Frontal Inf Orb R | ORBinf.R | 41, 32, -12 | Temporal Pole Sup R | TPOsup.R | 48, 15, -17 |
| Frontal Sup Medial L | SFGmed.L | -5, 49, 31 | Precentral L | PreCG.L | -39, -6, 51 |
| Frontal Sup Medial R | SFGmed.R | 9, 51, 30 | Rectus L | REC.L | -5, 37, -18 |
| Amygdala L | AMYG.L | -24, 0, -17 | Rectus R | REC.R | 8, 36, -18 |

Modular structures in the PL group were aligned with intrinsic brain subnetworks. Specifically, the PL-C1 module covered core regions in the FPN, including middle frontal gyrus (MFG), opercular part of inferieor frontal gyrus (IFGoperc), triangular part of Inferior frontal gyrus (IFGtriang), and inferior parietal (IPL) (Marek and Dosenbach 2018; Oliver et al. 2019). The PL-C2 module coincided with DMN (Buckner et al. 2008; Oliver et al. 2019). Contrary to the PL, we identified integrations of the FPN and the DMN in the OT group. In OT-C1 and OT-C2, the components of these modules did not show notable overlapping with the FPN or the DMN, and some components of OT-C3 belonged to the FPN and the DMN.

### Clustering coefficient and Average shortest path

We applied the Wilcoxon rank sum test to compare the clustering coefficients between the PL and OT groups. Among 90 brain nodes, 9 regions showed significantly smaller clustering coefficients in the OT group compared to the PL group *(p <* 0.01), including 5 frontal regions (left and right MFG, right ORBmid, right IFGoperc and right IFGtriang). We found a large proportion of nodes in the frontal network revealed relatively strong OT induced effects, compared to other brain areas. Moreover, an additional node, namely left IFGoperc, in OT showed significantly lower CC(*p* < 0.05). As suggested above, we focused on these six regions (5 frontal regions and left IFGoperc, including MFG.L, MFG.R, ORBmid.R, IFGoperc.R, IFGtriang.R and IFGoperc.L) in the following analysis. The complete results were shown in **Figure 3A**.

**Figure 3:**
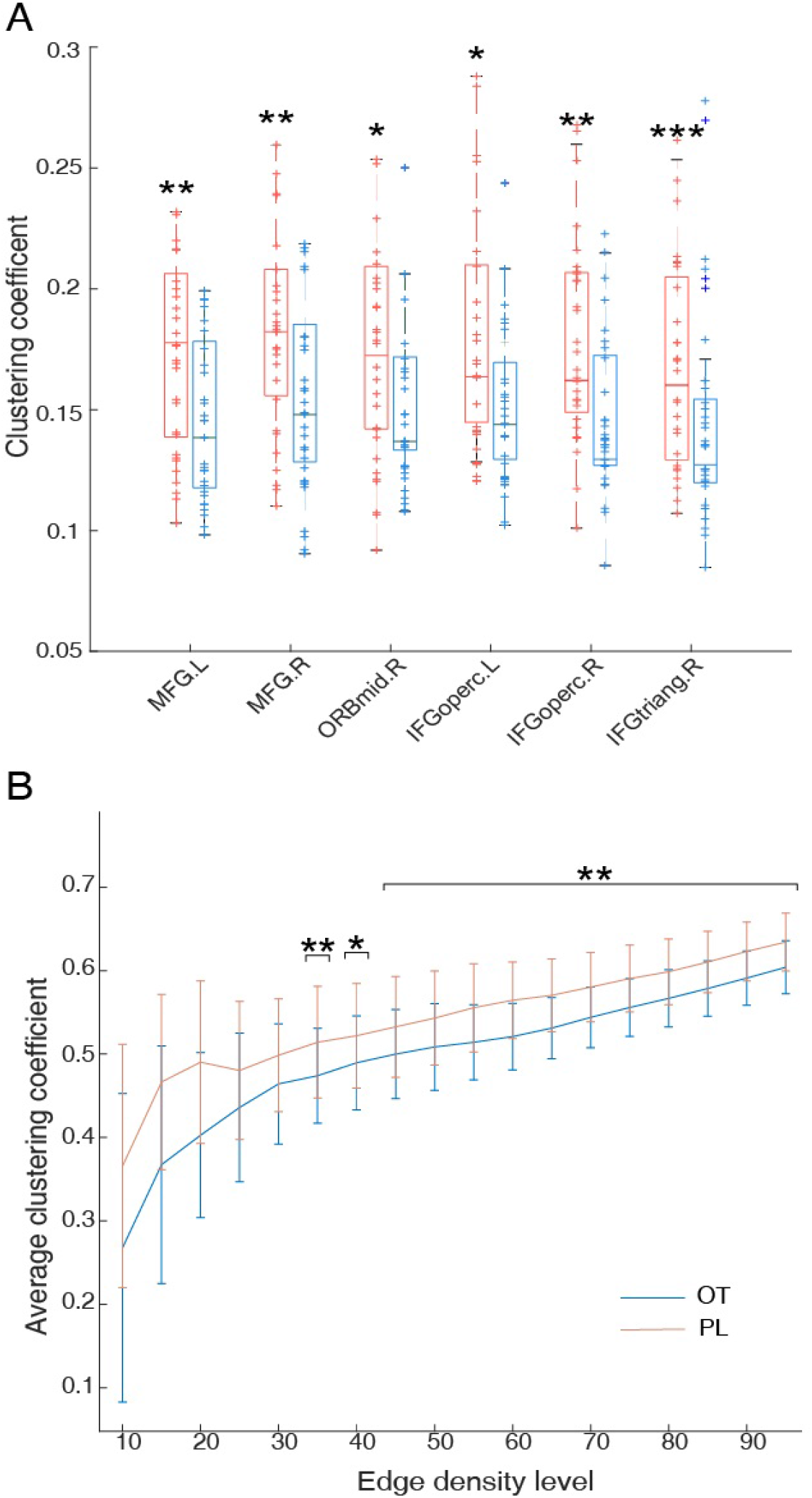
(A) Boxplot of the CC between OT (blue) and PL (red) groups in 6 frontal nodes. Dots represent each of two groups(29 and 28) individual CC. (B) Average CC between two groups with varying edge density. Error bar indicates a standard deviation. OT group showed smaller CC compared to PL. (* *p* < 0.05; ** *p* < 0.01; *** *p* < 0.001)

As listed in **Table 1**, nodes indicated with a star are the ROIs where clustering coefficients are significantly distinct between OT and PL groups, with smaller clustering coefficient values in the OT group. Notice that all these 6 nodes belonged to the FPN. In the OT group, the significantly smaller clustering coefficient values and larger average shortest path mentioned above could be viewed as a decrease in the small-worldness property within the FPN.

To avoid the influence of spurious correlation in fMRI data, we calculated the average CC of 6 frontal nodes for each group in different edge-density levels and applied the Wilcoxon rank sum test. In **Figure 3B**, we showed significantly larger CC in the PL group than the OT group (p < 0.01) at high edge-density level.

**Figure 4** illustrated the average shortest path in each group. All 6 frontal nodes in the OT group exhibited statistically higher ASP than the PL group.

**Figure 4:**
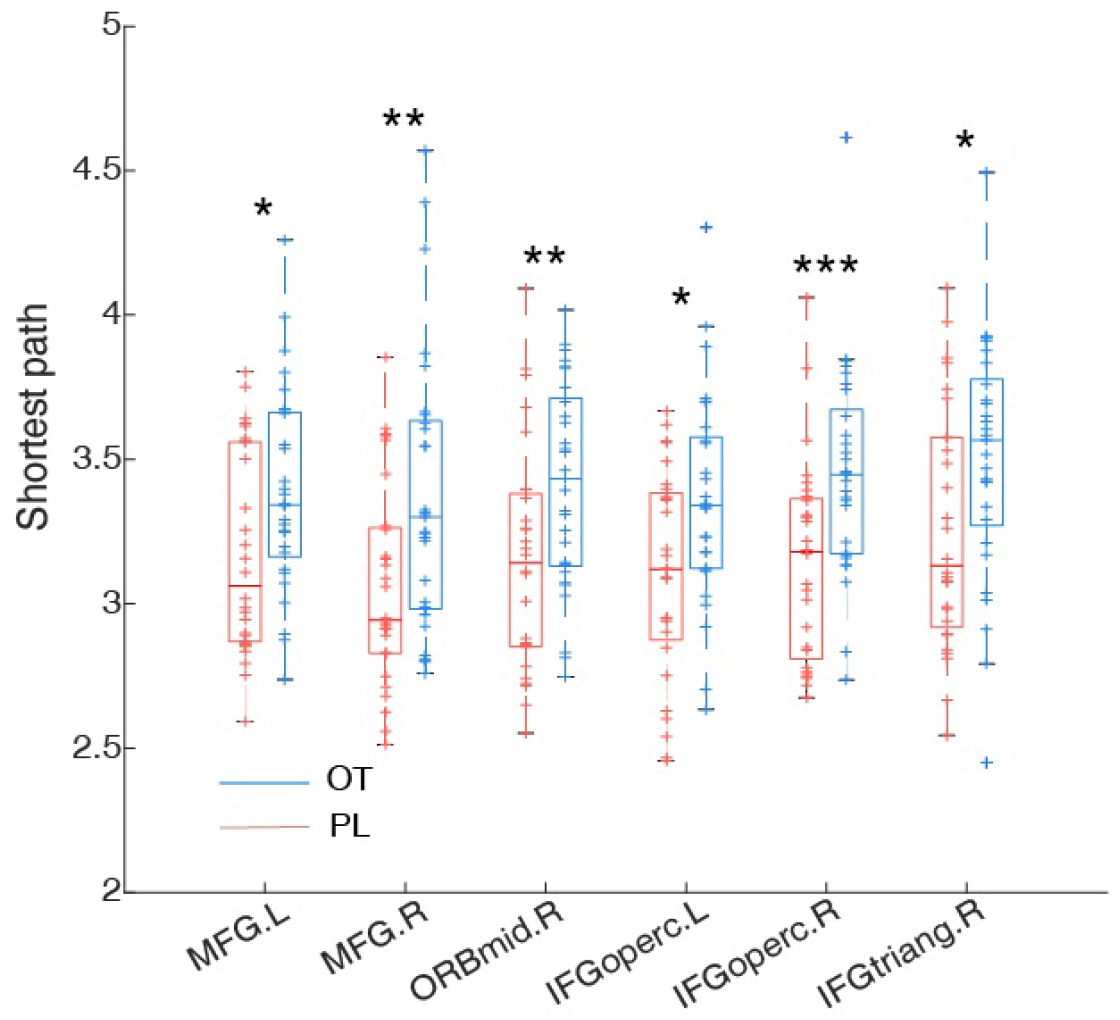
Average shortest path of the six frontal nodes. The OT group in blue and the PL group in red. (* *p* < 0.05; ** *p* < 0.01; *** *p* < 0.001)

Although the two groups have distinct CC and ASP values, both groups showed a significantly negative correlation between ASP and CC (See **Supplementary Table S1** for details). Furthermore, we performed eigenvector centrality and degree analysis. In the corresponding results, the calculated values of the two groups are closed, suggesting strong similarity. (See **Supplementary Figure S1 and S2**).

### Association between FPN and IRI traits

Statistical analysis (ANOVA) confirmed IRI performance uniformity in two groups. There is no significant difference between two groups in all three subscales (See Table S2 for details). To examine the relationship between network properties and personal traits, we performed a correlation analysis between CC and IRI subscales. **Figure 5** summarizes the CC-IRI correlation with networks at 3 varying edge-density levels. The OT group showed a significantly positive correlation between the fantasy score and CC, while the PL group showed a significant negative correlation. No other subscales showed significant correlation with CC. The detailed correlation between CC and fantasy were illustrated in **Figure 6**. Individuals with smaller fantasy values showed a more severe decrease of CC in the FPN for the OT group, implying that OT induced more substantial network alterations in the FPN.

**Figure 5:**
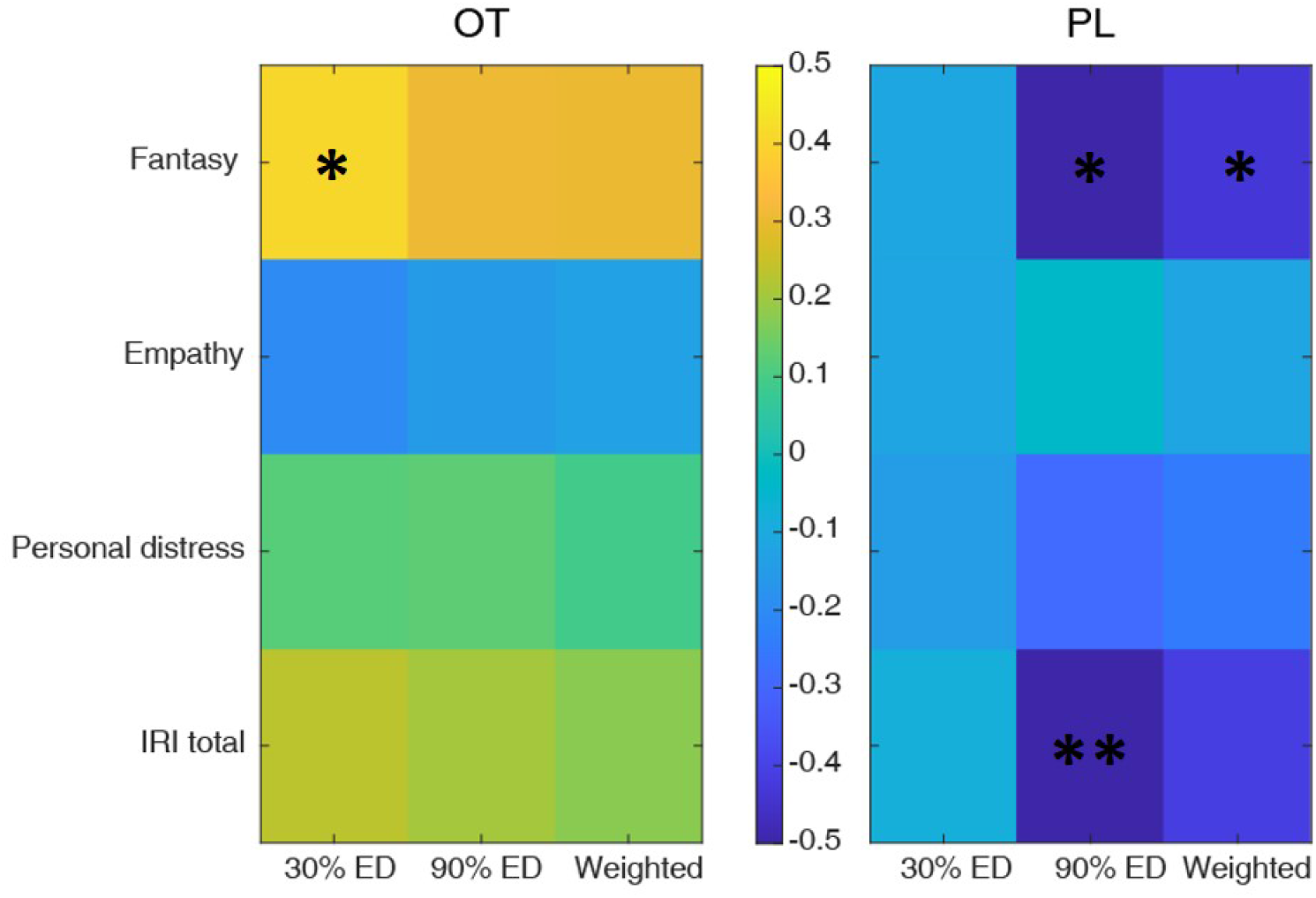
Correlation between average CC over 6 selected regions and IRI scale. Three different networks were used to calculate CC. (* *p* <0.05; ** *p* < 0.01; *** *p* < 0.001)

**Figure 6:**
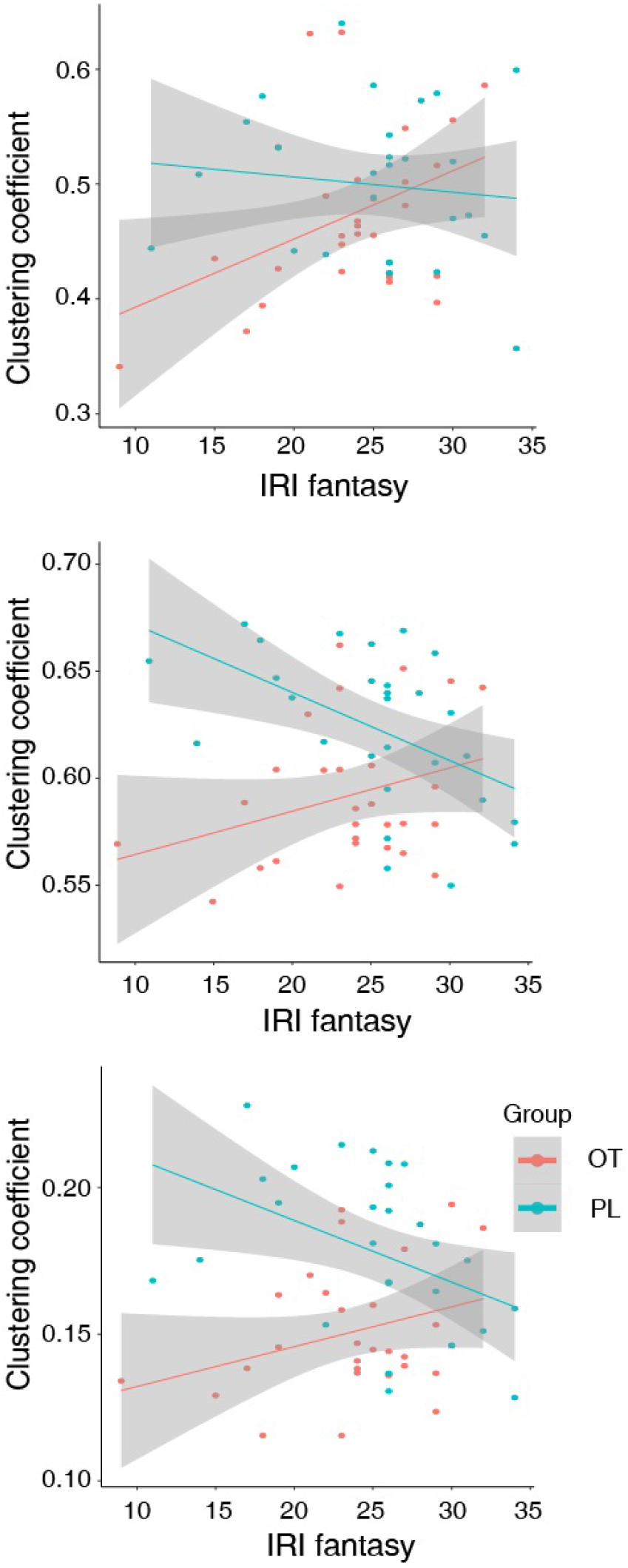
Scatter plot of individual IRI fantasy and CC with linear fitting. From left to right: 30%, 60%edge density level binary network and weighted network.

## Discussion

As an exploratory study, the objective of this work is to elucidate OT effects on frontal network characteristics. We hypothesized that OT elevates integration between subnetworks and reduces small-worldness within FPN. To test these hypotheses, we adopted measures including the community detection (CD), clustering coefficient (CC) and average shortest path (ASP). CD analysis showed less discriminative structures of FPN and DMN as well as increasing integrations between FPN, DMN and the limbic system in the OT group. CC and ASP analyses showed a disruption of small-worldness within FPN. In addition, we reported a negative correlation between CC and IRI-fantasy in the PL group but a positive correlation in the OT group. These findings are further discussed hereafter. In supplementary materials, we showed similar results of eigenvector centrality and degree distribution between two groups. Oxytocin may not have global topological effects (e.g., degree distribution) in the brain network. Eigenvector centrality is unable to reflect oxytocin effects in the frontal region.

### Structural alterations in frontal network

Consistent with previous studies (Eckstein et al. 2017; Sripada et al. 2013), we identified increasing connectivity between the limbic system (AMYG, HIP, and PHG) and the medial prefrontal cortex, supporting emotion processing and regulation. The anti-correlation between FPN and DMN has been found in previous research (Kelly et al. 2008). In general, the FPN and the DMN have opposite functions (e.g. mind-wandering and goal-directed). Therefore, the anti-correlation was thought to be crucial in adapting environment flexibly. Disruption of the anti-correlation was identified in patients with clinical symptoms like schizophrenia and obsessivecompulsive disorder (Jia et al. 2020; Stern et al. 2012). These studies provide evidence that increasing integration of the FPN and the DMN in the OT group may lead to alterations in cognition. Recent studies have identified two subnetworks within the frontoparietal control network. One subnetwork is closely related to the DMN and contributes to the introspective process. The other subnetwork is connected to the dorsal attention network and involved in working memory (Dixon et al. 2018; Murphy et al. 2020). A detailed examination of oxytocin’s effects in subnetworks of the FPN is needed in future research. Further analysis suggested that OT causes disruption of the small-worldness within the FPN. OT modules exhibit smaller modular size and a segregation pattern compared to PL modules. However, this does not guarantee OT modules to be more small-worldness. On the contrary, in OT modules, 6 nodes show significantly lower CC, compared to their states in the PL-C1 module (FPN). Additionally, OT modules show less distinguishable structures of the FPN and the DMN, as reflected in **Table 1**. Core regions of the FPN include IFGoperc L&R and IFGtriang.R showed significantly lower clustering coefficient in the OT group, compared to the PL group. In previous research, IFGoperc L&R and IFGtriang.R were identified as regions with functions in voice recognition (Elmer 2016; Schremm et al. 2018). In our study, IFGtriang decouples with the FPN and forms a community with significantly lower clustering coefficient. When exposed to infant crying, oxytocin has found to increase activation of IFGtriang (Riem, Bakermans-Kranenburg, Pieper, et al. 2011). Our research may provide a network-level explanation for such increased social signal sensitivity during OT treatment.

It is widely accepted that modular structures in the biological network represent functional units (Girvan and Newman 2002). In the brain network model, segregated modules correspond to dissociable cognitive components (Bertolero et al. 2015; Xu et al. 2017). One critical property of the FPN is that it serves as a flexible hub for cognitive control, which highly interacts with other networks and provides a functional backbone for prompt modulation of other brain networks (Marek and Dosenbach 2018; Zanto and Gazzaley 2013). Under OT administration, whether FPN still works as a backbone to modulate other subnetworks remain unknown.

### IRI Fantasy and brain network

Our study shows a significant correlation between individual average CC over 6 nodes in the FPN and the IRI fantasy scale, as seen in **Figure 6**, which is in line with previous studies (Golchert et al. 2017; Shi et al. 2018). For example, one previous study found more frequent reports of spontaneous mind-wandering were associated with an increasing integration between the FPN, DMN and limbic system (Golchert et al. 2017). Another study found weaker connectivity within the FPN is associated with less creativity in resting state (Shi et al. 2018). The smaller CC values in the OT group in our findings, indicating disruption of small-worldness within the FPN (**Figure 6**), and it may thus blur the boundary between FPN and other subnetworks. As it is critical to vicariously experience and to understand the affect of other people through imagination in social cognition, higher fantasy value implies individuals are more likely to put themselves in fictional situations. Thus fantasy value have potential associations with studies mentioned above. For instance, individuals with Autism Spectrum Disorder showed deficits in imaginative play as scarcity of pretend play a social activity (Wolfberg et al. 2012), fMRI study found that individuals with ASD had reduced neural responses in AI and ACC when they viewed other people in pain (Y.-T. Fan et al. 2014). There is also evidence showing atypical network distribution in ASD (Keown et al. 2017). The negative correlation between CC and IRI fantasy in the PL group aligned with previous research (Golchert et al. 2017; Shi et al. 2018). It is possible to find other correlations between personality traits and network property. Moreover, a critical issue is to understand the biological basis behind this diverted relationship of CC and IRI fantasy scale in OT and PL groups.

### Limitation and Future work

Several limitations in this study should also be mentioned. First, we recruited young, male, college students. However, the effect of OT on brain networks and their properties may vary with age, sex and education level. Whether our findings can be generalized to other groups of people remains unknown. However, it is also worth noticing that the graph theoretical analysis on fMRI signals used in this paper may be naturally applied to other groups. It is long recognized that the small sample size may reduce the reproducibility of fMRI studies. The sample size in our research is limited, which may affect the reproducibility of our study. The reliability of functional connectivity is becoming a concern of cognitive scientists. According to a recent meta-analysis (Noble et al. 2019), frontal and default mode networks are most reliable comparing to edges in other regions. Therefore, we may overvalue these networks in terms of oxytocin’s effects. Acquisition and processing strategies (Zuo et al. 2019), participants’ mental state could also influence reliability. To improve reliability, one essential approach is to collect high-quality data, reducing artifacts signal. It is also important to adopt optimized analytical strategies, which reduces the minimum data to achieved required reliability (Zuo et al. 2019).

Another limitation is that we did not examine temporal dynamic information for the network model in the current analyses. As indicated in previous research, we did not collect resting fMRI data before and after the OT administration (Wu et al. 2020). Additionally, we did not build a temporal dynamic network model. Our present study focused on the static network alterations after OT administration, which are limited in providing insights into causal relations of OT modulation effects. According to recent findings, functional brain network connectivity is dynamic even on the time scale of tens of seconds (Allen et al. 2014; Calhoun et al. 2014; Vidaurre et al. 2017; Yu et al. 2018), which emphasized the intrinsic slowly varying property of a large-scale fMRI-based brain network model. Indeed, on a network level, the study of the effects of OT remain in the early stages. Future work should incorporate temporal network properties into the model to pave the way for revealing causal relations in subnetwork alterations.

Traditional network models have limited insight regarding the rules underlying the transformation and encoding of neural response patterns in both local and distributed circuits (Sporns 2014). Concerning network controllability, a recent study reported that common graph metrics like the degree, path length, clustering coefficient, and modularity were not significantly different between mTBI patients and healthy people (Gu, Betzel, et al. 2017). In our study, although we observe less discriminative structures of the DMN and the FPN (which support cognitive control), this result does not lead to the conclusion about controllability in the DMN and the FPN. In essence, network measures used in our study are derived from network connectivity, while controllability is a novel idea rooted in the concept of network energy (Gu, Pasqualetti, et al. 2015; Pasqualetti et al. 2014), which may shed light to future network neuroscience research.

Together, this study investigated the modulation effects of OT on the frontal network based on graph theoretical analysis. We found substantial effects of OT on elevating the integration between the FPN, DMN and limbic system as well as disrupting small-worldness within the FPN. We also reported a strong association between FPN property and IRI fantasy score. The graph theoretical analysis on brain networks can be generalized to other applications, and provides a potential tool to assess the drug-induced changes on a brain network level.

