## Supplementary Figure S1, S2... for "Graph theoretic analysis reveals intranasal oxytocin induced network changes over frontal regions"

### Supplementary Material

November 9, 2020

#### 1 Eigenvector Centrality over frontal network

In **Figure S1**, we presented the results from 2 nodes as examples to demonstrate that the eigenvector centrality of frontoparietal network nodes in the PL group were significantly higher than the OT group ($p < 0.05$) at low-edge density, while they were not significant at the high-density level.

#### 2 Degree distribution

As shown in **Figure S2**, we plotted the degree distribution and fitted it with exponentially truncated power law ($p(x) = Ax^{-a}e^{-bx}$) [1]. The goodness of fit are $R^2 = 0.9762$ for OT; $R^2 = 0.9761$ for PL. The Jensen-Shannon Divergence of two discrete distributions is 0.0797, indicating large similarity between degree distributions of two groups.

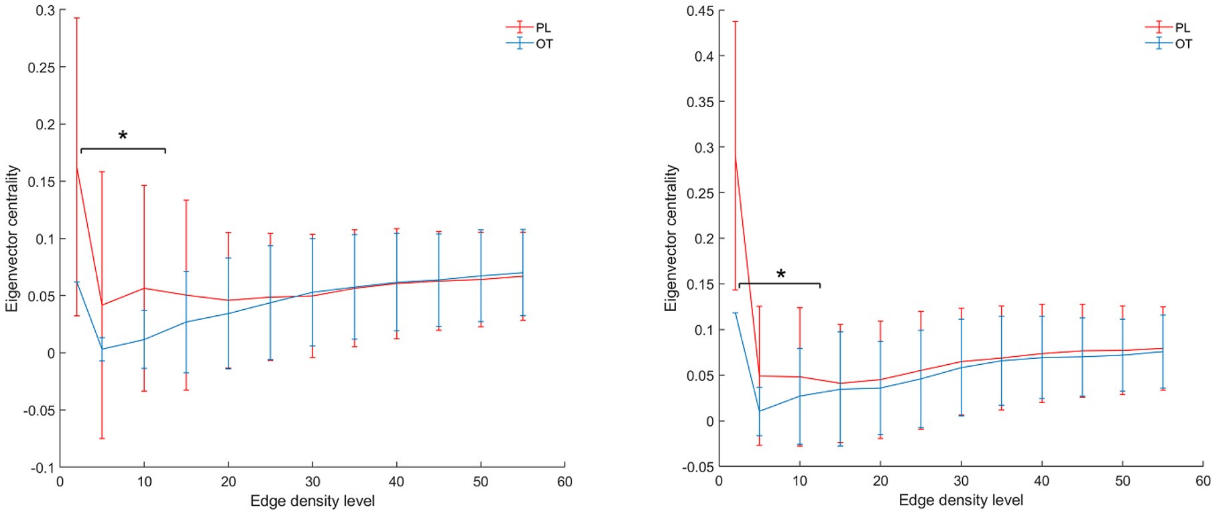


Figure S1: Eigenvector centrality between OT and PL across differnt edge density in MFG and r.ORBmid.Each plot is taken from a specific node in two groups. With large standard deviation and fluctuation, EC values in edge density level is not robust. While in high edge density level, values of OT and PL groups converge, showing similar pattern. (* $p < 0.05$; ** $p < 0.01$; *** $p < 0.001$)

Table S1: Correlation of CC and ASP in 3 regions

| | **OT** | | | **PL** | | |
|---|---|---|---|---|---|---|
| | Pearson's r | t | p-value | Pearson's r | t | p-value |
| MFG.L | -0.866 | -8.9918 | <0.001 | -0.914 | -11.499 | <0.001 |
| MFG.R | -0.924 | -12.537 | <0.001 | -0.889 | -9.987 | <0.001 |
| ORBmid.R | -0.821 | -7.460 | <0.001 | -0.920 | -11.965 | <0.001 |

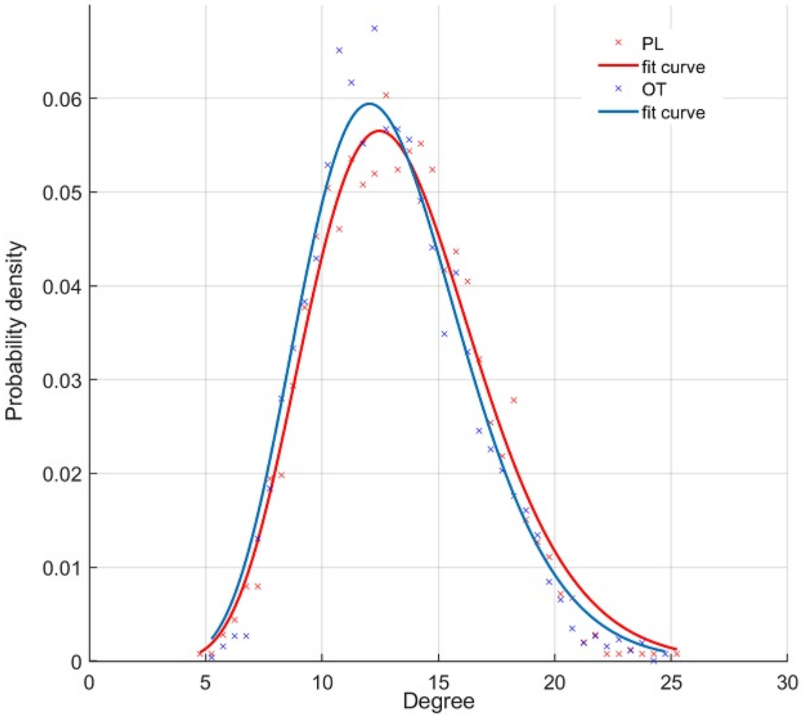


Figure S2: Degree distribution in OT and PL group.Two groups show similar patterns in degree distribution over the whole brain network model.

Table S2: Demographic and Clinical Characteristics

| | **OT group** | **PL group** | **t-score**[a] | **p-value** |
|---|---|---|---|---|
| age (year) | 22.86 (1.57) | 22.79 (2.27) | 0.134 | 0.894 |
| body weight (kg) | 65.87 (9.16) | 68.21 (9.37) | -0.970 | 0.336 |
| IRI total scores | 93.83 (10.81) | 95.10 (9.09) | -0.478 | 0.634 |
| Perspective taking[b] | 22.79 (3.60) | 22.03 (2.91) | 0.883 | 0.381 |
| Fantasy[b] | 23.14 (5.83) | 25.17 (5.06) | -1.419 | 0.161 |
| Empathic concern[b] | 25.83 (4.54) | 26.21 (4.16) | -0.332 | 0.741 |
| Personal distress[b] | 22.07 (3.94) | 21.69 (4.40) | 0.346 | 0.731 |

Data are expressed as mean (SD; standard deviation); n=30 for oxytocin group; n=29 for placebo group. Abbreviations: OT, oxytocin; PL, placebo; IRI, Interpersonal Reactivity Index.
a: t-ttest, two tailed.
b: Data missing from one OT group (n=29).

### 3 Psychological scales

Interpersonal Reactivity Index (IRI) is a widely-used, multidimensional assessment of cognitive reaction [2]. There are four IRI subscales, including perspective taking (PT), fantasy (FS), empathic concern (EC), and personal distress (PD). Each subscale includes seven questions. EC measures individuals' feelings of compassion and concern for others. FS describes the tendencies that respondents transpose themselves into fictional characters. PD indicates the extent that individuals feel uneasiness when exposed to the negative experiences of others. PT assesses unplanned attempts to adopt others' points of view [3].

Participants also took Neuroticism, Extraversion, Openness, Tas-20, and other scales. However, we did not use these scales in finding an association between personality traits and network property.

In Table S2, we showed results of statistical tests [4].

### 4 Calculation of IRI scores

The participant needs to answer 28 questions in total, seven questions in each subscale. In each question, participants need to quantify how a statement describes themselves, from 1 (poorly describe) to 5 (describe well). Researchers add up values answered by the participant in each subscale. Four total values can be obtained. In each dimension, a larger total value reflects a better ability in interpersonal reaction within that dimension.
